# A left-lateralized dorsolateral prefrontal network for naming

**DOI:** 10.1101/2024.05.15.594403

**Authors:** Leyao Yu, Patricia Dugan, Werner Doyle, Orrin Devinsky, Daniel Friedman, Adeen Flinker

## Abstract

The ability to connect the form and meaning of a concept, known as word retrieval, is fundamental to human communication. While various input modalities could lead to identical word retrieval, the exact neural dynamics supporting this process relevant to daily auditory discourse remain poorly understood. Here, we recorded neurosurgical electrocorticographic (ECoG) data from 48 patients and dissociated two key language networks that highly overlap in time and space critical for word retrieval. Using unsupervised temporal clustering techniques, we found a semantic processing network located in the middle and inferior frontal gyri. This network was distinct from an articulatory planning network in the inferior frontal and precentral gyri, which was invariant to input modalities. Functionally, we confirmed that the semantic processing network encodes word surprisal during sentence perception. These findings elucidate neurophysiological mechanisms underlying the processing of semantic auditory inputs ranging from passive language comprehension to conversational speech.

## Introduction

Word retrieval underpins human communication by accessing and retrieving word forms before articulation. Impairments in word retrieval (anomia) are very common following brain injury [1–3] and neurosurgical tissue removal [4–6]. Naming tasks, which seek to isolate word retrieval, are central to clinical assessment and basic research. Typically, a stimulus is presented visually or auditorily and the associated perceptual representations are then transformed into a shared pre-articulatory code [7, 8], which is in turn converted into an articulatory plan for overt word production [9, 10]. Clinical settings often employ a battery of naming tasks, commonly including visual naming, verb generation, reading, and repetition [9, 11–15]. These are widely used in intraoperative language mapping [11, 16–20], neuroimaging studies [21–25], neuropsychological assessments [14, 26, 27], and postsurgical outcome evaluation [13, 15, 28–31].

Two decades ago, Hamberger et al. observed a notable clinical gap: many patients with intact abilities to name visually presented stimuli experienced difficulties in word retrieval during everyday discourse. To address this, they introduced a task called auditory naming, where individuals were prompted to name items based on spoken descriptions [23, 32–35]. Intraoperative electrical stimulation mapping identified temporal cortex sites critical for auditory naming and predictive of post-operative naming abilities [11, 17]. Moreover, stimulation studies have revealed distinct functional segregation within lateral temporal regions, with specific areas, notably the anterior temporal lobe, supporting auditory naming, above and beyond visual naming [11, 16, 35]. Non-invasive neuroimaging studies sampling all of cortex have provided evidence for similar differential patterns, including both the anterior temporal lobe and prefrontal regions such as the Inferior Frontal Gyrus (IFG)[5, 22, 36]. However, all of these studies are inherently limited in their temporal resolution.

A growing body of literature has used intracranial Electrocorticography (ECoG) to study auditory naming. This technique, with a combined high spatial and temporal resolution, has revealed IFG activity during multiple overlapping time periods: immediately after stimulus onset [37, 38], at stimulus offset [37, 39], prior to response onset [39, 40], and throughout stimulus presentation [41]. This variability in activation pattern implicates IFG in supporting multiple cognitive processes throughout the auditory naming task. Early activation during auditory stimuli presentation suggests IFG plays a role in semantic processing, as the listener perceives the description evolving in time [37, 41]. Later, activation prior to response onset suggests its additional role in articulatory planning. This is in line with studies establishing IFG in articulatory planning during non-semantic tasks (i.e. visual word reading and auditory repetition) [10, 42]. While auditory naming requires both these cognitive processes, the neural activity subserving them overlaps spatially and temporally. In order to dissociate these overlapping neural responses, it is necessary to closely contrast auditory naming tasks with matched articulatory and non-semantic controls. However, studies to date have focused on semantic processes alone (i.e visual and auditory naming), obscuring a dissociation between semantic and articulatory processing.

To address this, we designed an experiment that controls for modality, semantics, and articulatory target. By incorporating four distinct tasks that elicit the same spoken word through different combinations of cognitive pathways, we predicted that overlapping functions, such as articulatory planning, and modality-specific retrieval, are dissociable with a data-driven method. Using ECoG data from a large cohort of neurosurgical patients (N=48), we identified two distinct spatiotemporal networks that separate articulatory planning from semantic processing along a ventral-dorsal gradient. Spatially, semantic processing spanned dorsal frontal cortices, while articulatory planning was restricted ventrally. Further, we validated that the dorsal network encodes increased semantic load during sentence presentation and showed that this network is left-lateralized. These findings constitute novel evidence for a hub in dorsal IFG and MFG responsible for semantic integration, a critical linguistic function for behaviors ranging from reading to discourse.

## Results

In order to investigate the spatiotemporal dynamics leading up to lexical retrieval, we recorded ECoG signals from 29 (25 left grid, 4 bilateral) participants sampling the left hemisphere across four tasks: visual naming (VN), visual word reading (VWR), auditory naming (AN), and auditory word repetition (AWR). The tasks were designed to produce the same set of words while varying the routes of retrieval, controlling for task modality and semantic access (Fig. 1A, see Methods: Experiment Setup). Neural responses were quantified using high gamma broadband (70-150 Hz) (See Methods: Data Acquisition and Preprocessing), a widely applied marker of local cortical activity that correlates with underlying neuronal spike rates and the BOLD response [43]. We first report single-trial activity across task-active electrodes locked to articulation in three regions of interest: IFG, Middle Frontal Gyrus (MFG), and precentral gyrus (Methods: Task-active Electrode Selection). Temporally, we found enhanced responses before articulation across all three regions (Fig. 1B) which exhibited specificity for the semantic tasks compared with control. (one-way *t*-test: VN vs. VWR: IFG, *p <*1e-5, *t* = 7.96; MFG, *p <*1e-5, *t* = 20.86; precentral, *p <*1e-5, *t* = 9.19; AN vs. AWR: IFG, *p <*1e-5, *t* = 16.50; MFG, *p <*1e-5, *t* = 25.39; precentral, *p <*1e-5, *t* = 7.36, see Methods: Statistical Tests). We confirmed this pattern by focusing on a pre-articulatory window (-500ms - 0ms) within each electrode, showing enhanced responses for the semantic tasks with maximal activation in auditory naming (Fig. 1C). The results provide evidence for the highest neural recruitment in prefrontal cortices during auditory naming.

**Fig. 1.**
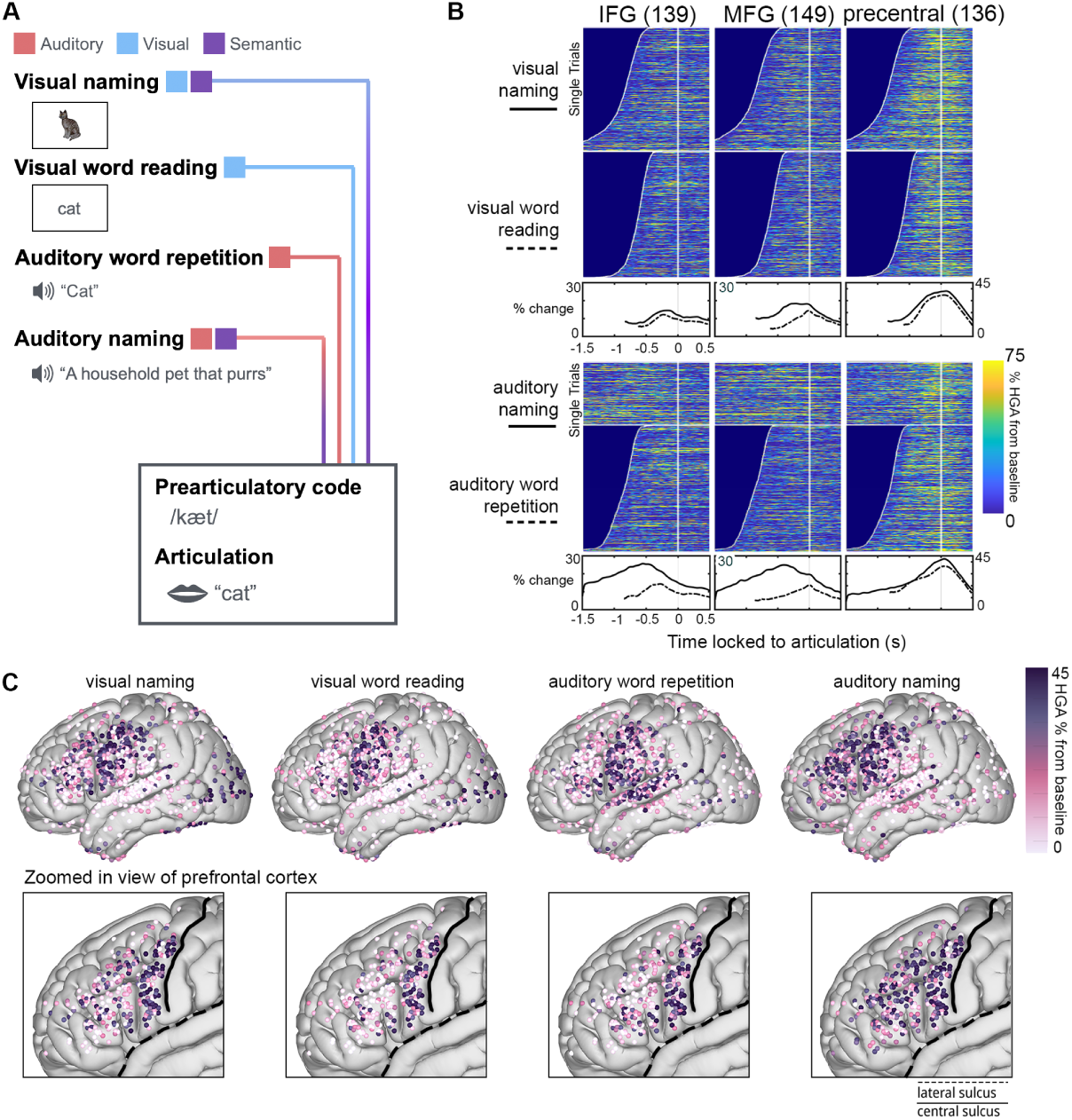
(A) Experimental paradigm depicting the four tasks underlying varying routes of retrieval and cognitive processes leading up to articulation. (B) Vertically stacked single trials sorted by response time are shown for tasks locked to articulation. The white line indicates stimulus onset. Single-trial high gamma activity (HGA) peaked before articulation in MFG and IFG and peaked at articulation in precentral gyrus. Group average across trials is shown for naming tasks (top: visual naming; bottom: auditory naming) in solid black lines and their controls in dash lines (top: visual word reading; bottom: auditory repetition). The cutoff was determined by median response time within each task. (C) Top: Spatial dynamics of HGA during a pre-articulatory window (-500ms - 0ms) for each task. Colors indicate average neural activity across electrodes. Only significant electrodes across perception and production are shown. Bottom: Enlarged view of the prefrontal cortex.

To statistically evaluate the semanticand modality-specific factors modulating neural activity before articulation, we employed a multilinear encoding model, which can predict high gamma activity from multiple features and their interactions (see Methods: Multilinear Factor Model). Our experiment was designed such that each task produces the same words and contains a unique combination of both feature dimensions (Fig. 2A). We first quantified the significance of each feature across taskactive electrodes in the pre-articulatory window (-500ms - 0ms). We found that the auditory modality mostly predicted activity in STG (Fig. 2B, top), the visual modality predicted activity in occipital cortex (see Supplementary Fig. S1) and auditory semantics strongly predicted activity across both IFG and MFG (Fig. 2B, middle, showing significant electrodes, FDR corrected *q* =1e-3). Strikingly, visual semantics dissociated from its auditory counterpart with a more posterior distribution overlapping with MFG (two-sample *t*-test, *t* = 3.94, *p <* 1e-3, see Fig. 2B, middle and bottom, and Supplementary Fig. S1, D-F). To ensure the effect represents sustained semantic access before articulation, we employed a temporal moving window approach (-750ms - 500ms). Sustained and significant (FDR corrected *q* =1e-3) modality-specific semantic effects were most prominent from -500 ms to -250 ms across frontal cortices and dissipated by the time of articulation (Fig. 2C). Taken together, these results provide evidence for a posterior to anterior gradient encoding semantic processing for the auditory and visual domains, respectively, necessary for word retrieval.

**Fig. 2.**
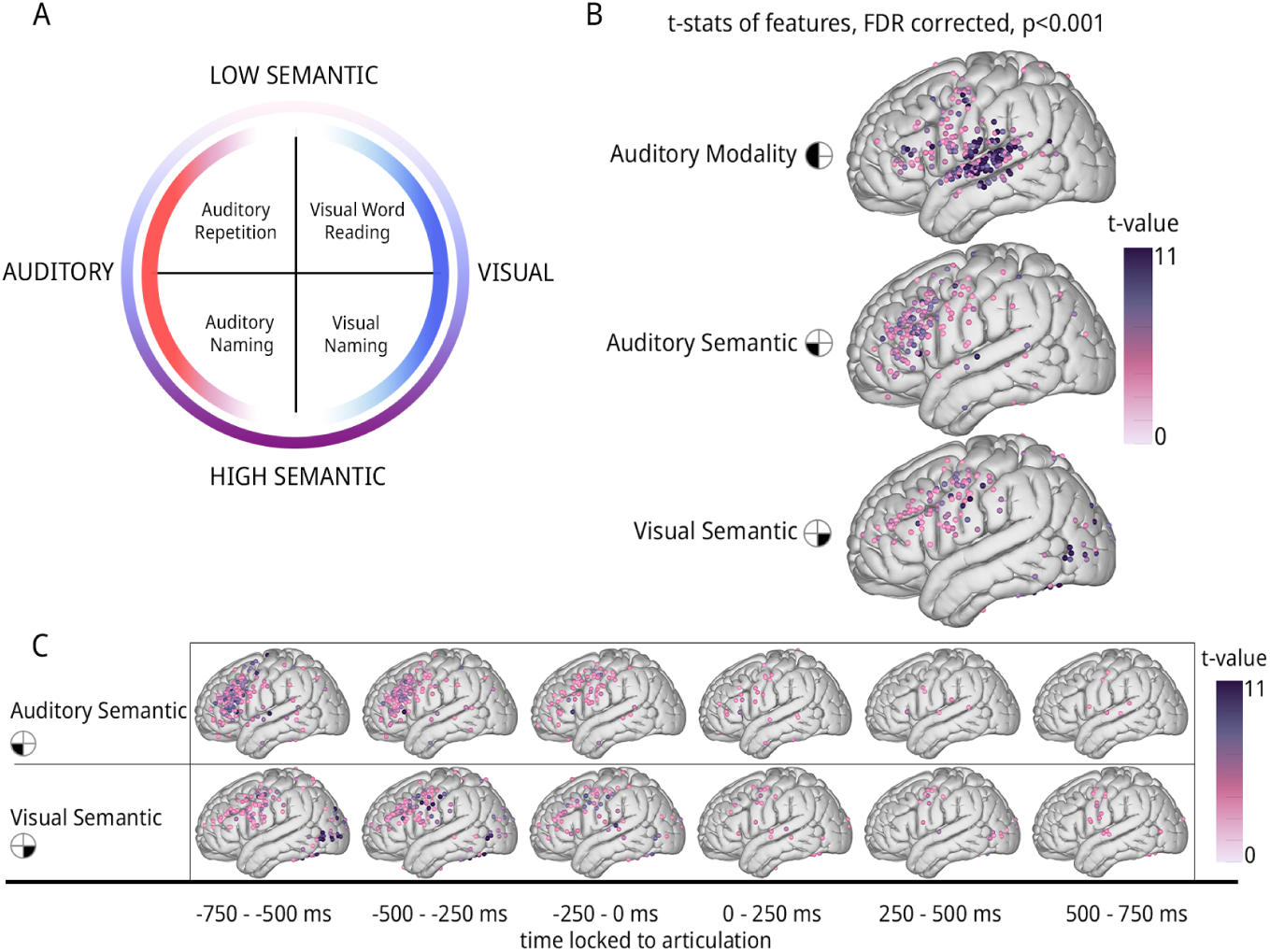
(A) Modality and semantic feature paradigm for constructing encoding model. (B) *t*-value map of encoding results -500 ms before articulation. Electrodes that significantly encode modality (top), auditory-specific semantics (middle), and visual-specific semantics (bottom) are projected on the brain. (C) Progression of visual- and auditory-specific semantic encoding map over time, locked to articulation onset.

Given that these results spanned multiple frontal regions (e.g. IFG, MFG, and precentral gyrus), we were interested in understanding the prototypical temporal profiles across frontal cortex. In contrast to previous studies reporting averaged responses within a region, we did not want to impose any anatomical constraints. Therefore, we chose Non-Negative Matrix Factorization (NMF, see Methods: Non-negative Matrix Factorization) as a data-driven clustering method to capture the shared pattern from neural activity across all tasks, cortical sites, and participants. This approach identified five networks with unique temporal profiles in-line with sensory, pre-articulatory, and motor functions. (Fig. 3A). Two sensory networks cleanly captured stimulus-induced sensory responses in auditory and visual cortices (see Fig. 3A, Supplementary Fig. S2C). Three major networks were found across frontal cortices (Fig. 3A,B). The first network peaked before articulation, exhibiting enhanced activity for auditory naming relative to the other tasks (*p <* 1e-5, Kruskal-Wallis test, see Methods: Statistical Tests), and was spatially distributed across IFG and MFG (Fig. 3B, top), similar to the distribution of auditory semantic information we found in Fig. 2B. The second network also peaked before articulation (mean peak timing = -158 ms, SEM = 16 ms); however, activity did not differ across tasks (*p* =0.173, Kruskal-Wallis test) and was spatially distributed across IFG and speech motor cortex (SMC, i.e. peri-rolandic). Lastly, the third network peaked after articulation (mean = 93 ms, SEM = 11 ms), was spatially distributed across SMC, and did not differ across tasks (*p* =0.70, Kruskal-Wallis test). It is noteworthy that a minority of electrodes across SMC (especially dorsal precentral gyrus) also exhibited stimulus related and naming responses (Supplementary Fig. S2B). In summary, our results demonstrate the existence of two spatially and functionally distinct cortical networks preceding speech within frontal cortex: one task-invariant, suggesting its role in speech planning, and the other naming-specific, suggesting a role in mapping from auditory to semantic representations.

**Fig. 3.**
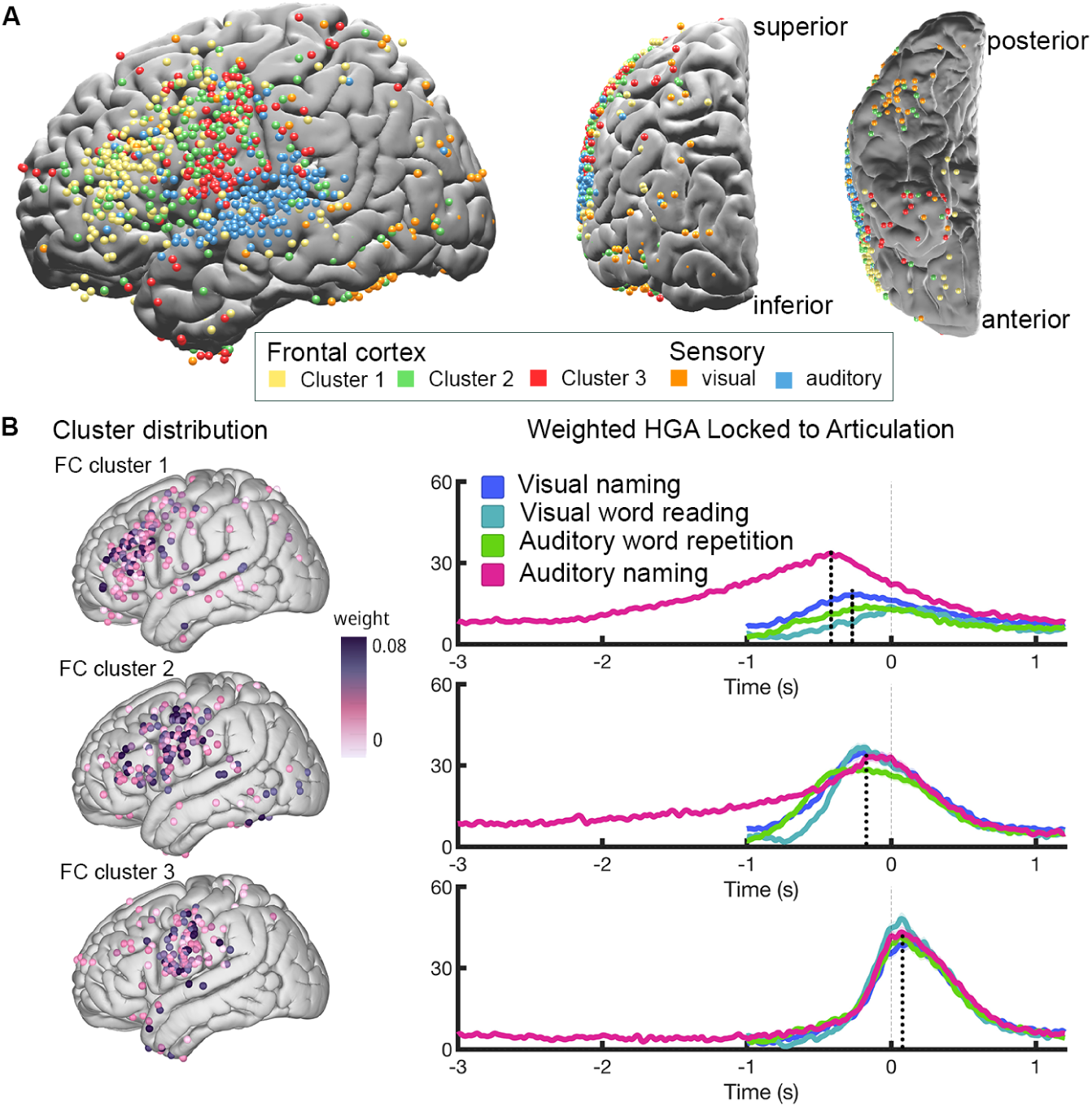
(A) Spatial positions of electrodes color-coded by NMF cluster membership. The desired number of clusters was identified with relative variance explained (See Supplementary Fig. S2A and Methods: Non-negative Matrix Factorization). (B) Left: the NMF weight distribution of electrodes in three frontal clusters. Right: the weighted average of neural high gamma activity across electrodes in the cluster, locked to articulation. Dashed lines mark the peak of the temporal profile.

We then investigated the extent to which the observed functional networks were part of the language network. Given that language production is predominantly leftlateralized, we expanded our analysis to include a cohort of patients with right hemisphere coverage (N=23, see Methods: Participants Information). We repeated the same clustering approach but using task-active electrodes from both hemispheres and statistically assessed the degree of hemispheric lateralization (See Methods: Permutation). This analysis replicated our left-hemisphere clustering results by identifying five networks with similar temporal profiles to the first left-hemisphere cluster analysis (see Supplementary Fig. S3B and S3C). The speech motor network exhibited a bilateral distribution (Fig. 4A, bottom) and the ratio of recruited task-active electrodes was not significantly different between the two hemispheres (*p* =0.1443, permutation test, see Methods: Permutation, Fig. 4B, bottom). In contrast, we found significant asymmetry in the task-active electrode ratio in both the naming-specific (*p <*1e-5, permutation test) and pre-articulatory networks (*p <*1e-5, permutation test) that were overwhelmingly left-lateralized. These results suggest that the left-lateralized naming-specific activity is likely crucial for language processing rather than serving as a domain-general component (e.g. working memory, which is bilateral [44]).

**Fig. 4.**
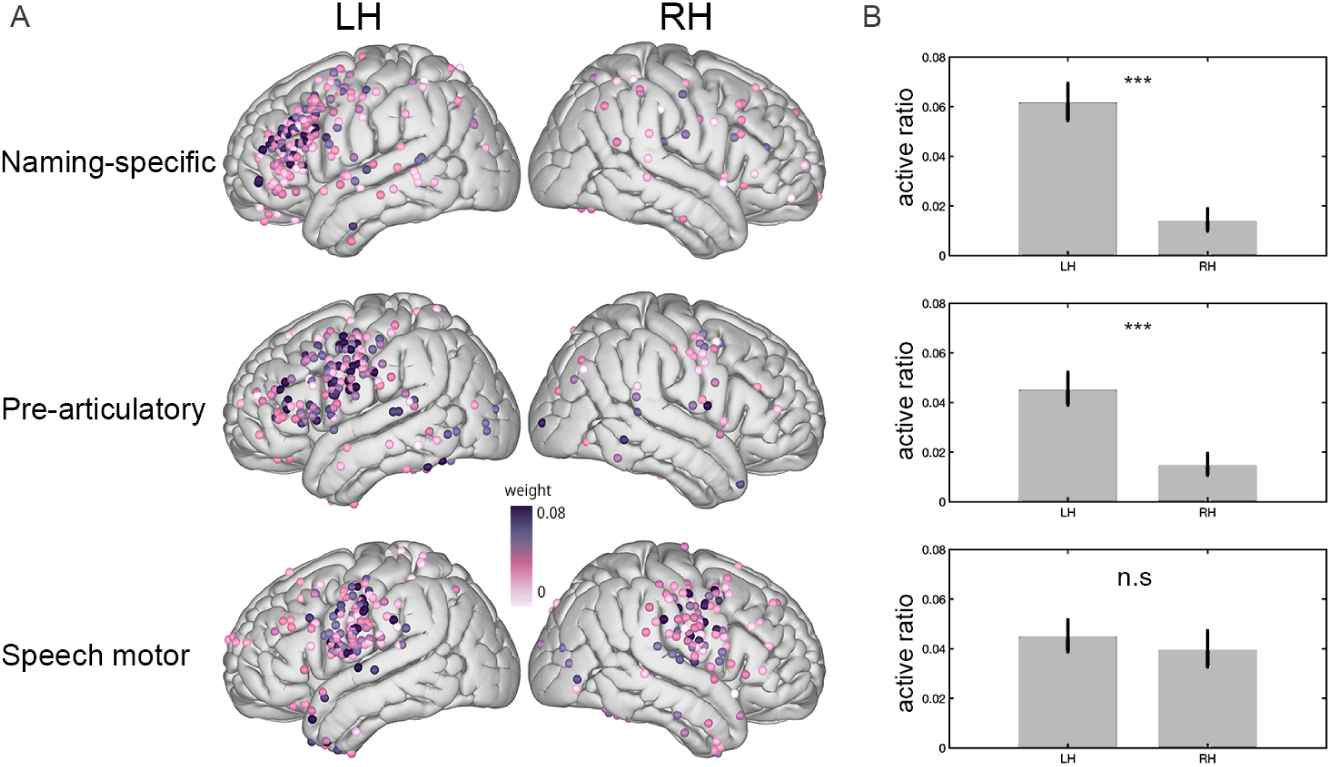
Clustering results in a cohort with bilateral coverage. The desired number of clusters (5) was identified with relative variance explained (See Supplementary Fig. S3A and Methods: Non-negative Matrix Factorization). (A) NMF weight distribution of electrodes in three frontal clusters. (B) Active ratios (task-active electrodes vs. all) for the left and right hemispheres are shown with bootstrapped confidence intervals depicted in the error bar Methods: Bootstrapping. Significance between left and right hemisphere active ratio was determined by a permutation test (see Methods: Permutation).

We were interested in understanding the underlying representation of these cortical networks. One popular approach in neuroscience to study neural representations is the use of encoding models. We employed such an approach using the multivariate temporal response function (mTRF; see Methods: Encoding Model), that predicts neural activity during auditory naming based on three features: acoustic information (i.e., speech envelope), task engagement (i.e., task structure), and word surprisal (i.e., the unexpectedness of a word within a given context; Fig. 5A). These features were selected to capture low-level sensory processing (acoustic), attentional demands (task engagement), and semantic integration (word surprisal) respectively (see Supplementary Fig. S4C for encoding results). To disentangle the individual contributions, we performed variance partitioning to quantify the unique variance explained by each feature, revealing the degree to which one feature explained variance not captured by the others (Methods: Variance Partitioning). In a representative STG electrode, we demonstrated the encoding variance for three raw models (acoustic, task engagement, and surprisal) and the unique variance explained by each representation, validated through a permutation test (Fig. 5A, right). Across cortex, significant word surprisal (Fig. 5B, top) was predominantly localized in IFG and MFG (left-lateralized) and STG (bilaterally). Significant acoustic representation was mainly localized in STG and dorsal precentral gyrus (bilaterally; Fig. 5B, middle). Representation of task engagement was mainly found in precentral gyrus (bilaterally; Fig. 5B, bottom). The left-lateralized spatial distribution of surprisal encoding was consistent with our previous clustering findings (Fig. 4A), and we quantified the surprisal encoding across these networks (Fig. 5C). Word surprisal was significantly greater for the naming-specific network compared with the other frontal networks (Wilcoxon rank sum test *p <*1e-5, see Methods: Statistical Tests. Speech motor: *z* = 8.91; pre-articulatory: *z* = 7.21; speech motor vs. pre-articulatory was not significant: *p* = 0.31. See Supplementary Fig. S4D and S4E for the encoding and unique variance explained results across all networks). These findings suggest that the naming-specific network mainly encodes word surprisal. Taken together, our results demonstrate that auditory semantic information, particularly concerning word surprisal, is encoded by a frontal network spanning IFG and MFG. This pattern is distinct from the pre-articulatory planning activity associated with speech and is left-lateralized.

**Fig. 5.**
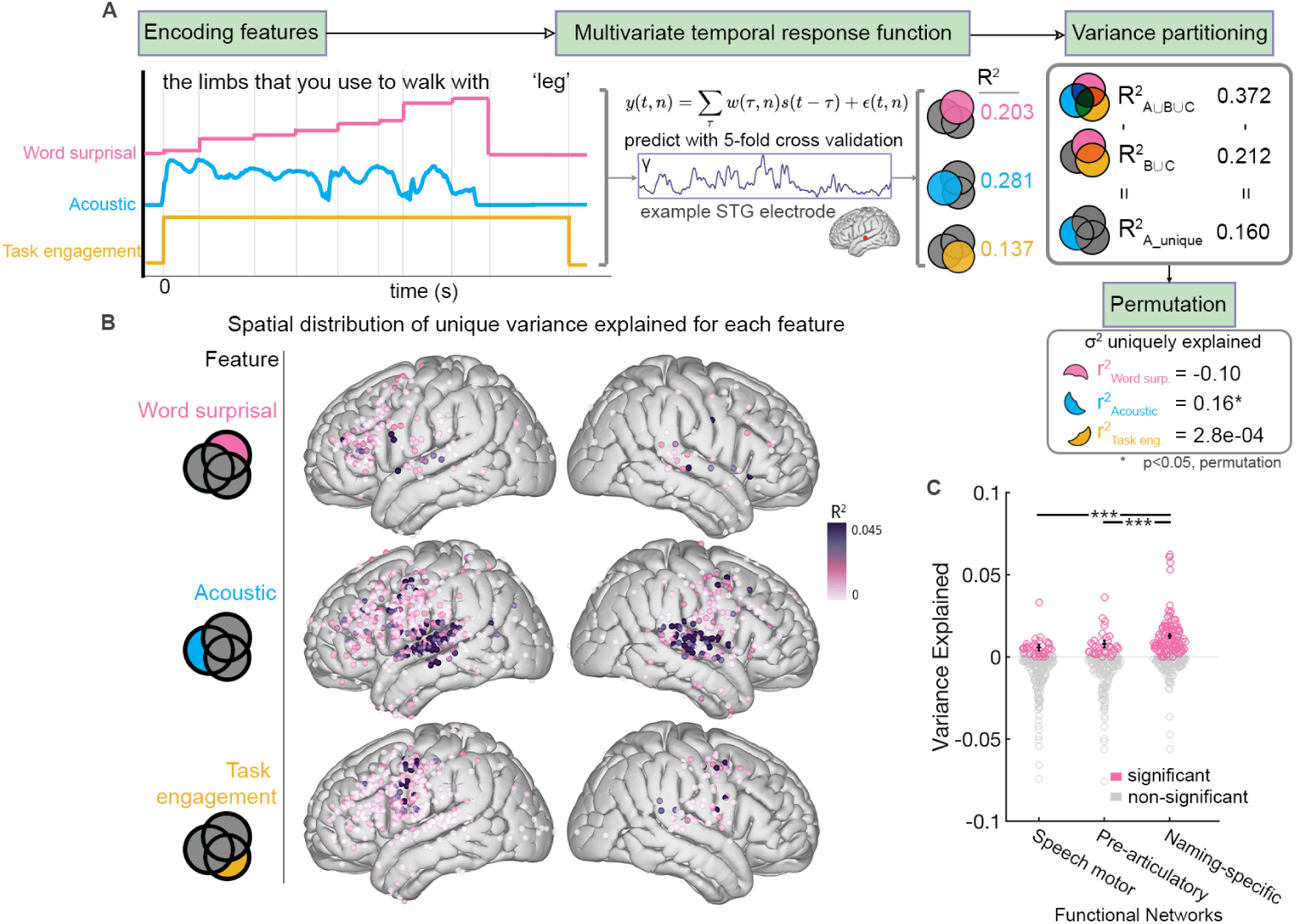
(A) Paradigm of encoding analysis. Encoding models were built based on representations of increment of word surprisal (top), acoustic (average spectra, middle), and task engagement (bottom). Unique variance was calculated by variance partitioning, and significance was tested by permutation. (B) Variance explained for each representation is depicted across cortex. (C) Significant surprisal model values determined by permutation (pink) are plotted for each network identified by the NMF analysis. Each dot represents the variance partition R^2^ of an electrode. A Wilcoxon rank sum test shows that the naming-specific network encodes more word surprisal information than the other two frontal networks.

## Discussion

Word retrieval is a fundamental human ability, critical for daily life communication and clinical assessment. Here, we recorded direct neurosurgical recordings to systematically probe lexical retrieval across clinical naming tasks. Our results highlight the spatial dissociation in the prefrontal cortex between auditory and visual naming. We then disentangled overlapping pre-speech neural activity supporting different functions into two frontal networks: one for auditory naming dorsally and the other for task-invariant articulatory planning ventrally. We further showed these networks are lateralized to the left hemisphere, implicating them in core language functions. Lastly, using supervised encoding models, we established that dorsal frontal activity best represented semantic integration of words. Taken together, these findings underscore an important role of dorsal prefrontal cortex in semantic integration and auditory-based lexical retrieval.

Our findings (Fig. 2) provide direct neurophysiological support for a dissociation between visual versus auditory naming across frontal cortices. Such a dissociation has been previously demonstrated in the temporal cortex alone and supported by stimulation studies rather than neurophysiological recordings [12, 45, 46]. Although previous intracranial neurophysiological studies have shown higher activation across the frontal cortex in auditory naming [37, 39, 41], they did not report a clear spatial dissociation. This is likely due to the task design which did not include non-semantic controls and matched articulatory targets. In particular, stimuli in several previous studies [37, 39] did not fully constrain the articulatory target (for example, “what is flying in the sky?”[37, 39, 47] in contrast to “a vehicle that flies through the air” see first item in Table S5: Auditory naming stimuli). Our study addressed this by ensuring that the same words were produced in each of the four tasks, allowing for a controlled comparison across modality-specific semantic processing (See Supplementary Fig. S1B for comparison without modality-specific control). Further, previous task effects were often assessed through mean activity comparison [48] or odds ratio [41] by averaging across electrodes within predefined anatomical regions. We tested the effect within each electrode, and showed for the first time, a gradient of modality-specific semantics activation along the anterior-posterior axis in the left prefrontal cortex (Fig. 2B, and Supplementary Fig. S1C for right hemisphere coverage). Across this gradient, auditory-semantics was encoded more anteriorly and visual-semantics more posteriorly. Our findings suggest a modality-specific modulation in the prefrontal cortex for semantic modalities, aligning with studies arguing for modality-specific attention networks in the prefrontal cortex [49–51].

Our study reveals that prefrontal neural activity subserves two distinct cognitive processes preceding speech (i.e. semantic and articulatory). Previous ECoG studies on naming employed a region of interest approach, which can obscure spatially overlapping activation patterns [37, 39–41]. Moreover, previous studies focused less on the shared temporal dynamics, as their analyses were based on task-active electrodes selected from the entire trial period [41]. Our study overcomes these issues by applying a clustering approach, which extracts shared temporal profiles without imposing anatomical constraints. Our reported ventral network is task-invariant and peaks before articulation. This is in line with the findings of IFG’s role in speech arrest [52–54] as well as pre-articulatory planning [10]. Recent studies have highlighted the important role of precentral gyrus in speech planning [55, 56], consistent with our results showing peak activation of precentral gyrus before articulation. Our dorsal network encompassing IFG and MFG showed naming-specific activity. This finding is in line with recent literature examining the role of MFG in speech and language [55, 57, 58]. Previous studies interpreted this enhanced MFG activity as higher working memory load [39], as bilateral dorsal lateral prefrontal cortex plays a role in working memory [44], and verbal auditory working memory during speech [59, 60]. However, our results showed a strong left-lateralization for this cluster (Fig. 4), implicating its role in higher-order language functions rather than working memory. Our findings are in line with Kambara et al., who showed that auditory working memory mainly resulted in enhanced precentral gyrus activity [40]. Taken together, our spatial dissociation between pre-articulatory and semantic networks (Fig. 3) suggests that a frontal semantic node might be missing in current language models that argue for a multi-stream process [61, 62].

Our study further probed the underlying representation of this naming-specific activity, and found evidence supporting its role in integrating word surprisal during sentence processing (Fig. 5). This finding is in line with literature showing IFG’s role in semantic and syntactic processing during sentence listening [63] and its word position sensitivity observed in serial visual word integration [64, 65]. However, our results suggest a more spatially distributed network that spans both IFG and MFG, rather than IFG alone (Supplementary Figure S2B). We employed mTRF models together with variance partitioning in order to control for dependencies between features [66]. This approach revealed the degree to which the word surprisal feature explained unique variance not captured by acoustic and task engagement features, thus making the results biologically interpretable. However, we acknowledge that this approach inherently produced negative values in some cases, as has been reported in prior studies [66–70]. Our permutation testing verified that these values were nonsignificant and therefore should be interpreted as noise, in line with previous studies that have ignored negative values [67, 69, 70]. Taken together, our data suggest that auditory semantic integration occurs across a distributed network spanning IFG and MFG, and is likely recruited during daily conversation [32].

The capacity to access words is critical for daily discourse communication and clinical assessment. Our findings demonstrate the recruitment of a dorsal prefrontal network specific for this function. Our unsupervised approach underscores that left-lateralized language functions are represented across cortical regions and are not limited to traditional boundaries. Furthermore, our encoding analyses suggest that semantic auditory integration is supported by activity in frontal sites, among others. Collectively, these results highlight a new perspective on the function of the dorsal prefrontal cortex in integrating information for communication.

### Limitations of the study

One limitation of our study lies in the inherent sampling bias of intracranial electrode coverage, which is based on clinical needs and was primarily concentrated over perisylvian cortices. This constraint is common in intracranial investigations but limits our ability to include other cortical regions in our analyses. Notably, the anterior temporal lobe (ATL), a region implicated in both naming and higher-order linguistic processes such as phrase- and sentence-level comprehension [5, 63, 71], was underrepresented in our dataset, potentially masking contributions from this critical area.

Our approach used data-driven clustering to dissociate cognitive processes without imposing anatomical constraints. However, we acknowledge that these results could potentially be modulated by the data provided to the algorithm. Future studies could include additional control tasks such as semantic categorization, or verb generation which could replicate our auditory-naming specific response in addition to clusters we could not examine with our set of tasks.

## STAR Methods

### EXPERIMENTAL MODEL AND STUDY PARTICIPANT DETAILS

#### HUMAN PARTICIPANTS

All experimental procedures were approved by the New York University School of Medicine Institutional Review Board. We used neurosurgical recordings from 48 patients (20 females, mean age: 28.0 ± 12.59 yo, 18 right grid, 25 left grid, 4 bilateral hemisphere coverage, and 1 stereo coverage. See Supplementary Table S1: Participant Demographics) undergoing neurosurgical evaluation for refractory epilepsy were included in the analysis. Patients implanted with subdural and depth electrodes provided informed consent to participate in the research protocol. Electrode implantation and location were guided solely by clinical requirements. We have 5 patients who consented separately for higher density clinical grid implantation, which provided denser sampling of the cortex. Surface reconstructions, electrode localization, and Montreal Neurological Institute coordinates were extracted by aligning a postoperative brain Magnetic Resonance Imaging (MRI) to the preoperative brain MRI using previously published methods [72]. We formally tested for a main effect of gender (with a random effect of patient) fitting a linear mixed effect model predicting average neural activity across tasks, and we found no significant result across time locked both to perception and production (*p >*0.05, FDR corrected for multiple comparisons)

### METHOD DETAILS

#### Experiment Setup

Participants were instructed to complete four tasks to produce the same target words in response to certain auditory or visual stimuli (Fig. 1A): visual naming (VN, overtly name a word based on drawings), visual word reading (VWR, overtly read the word presented on the screen), auditory naming (AN, overtly name a word based on auditorily presented description, and auditory word repetition (AWR, overtly repeat an auditorily presented word). The visual naming stimuli were selected from the normative dataset by Rossion and Pourtois [73], which is a colored adaptation of Snodgrass and Vanderwart’s original database of 260 black-and-white line drawings. For our study, we selected 50 stimuli, with their indices to the original dataset listed (Table S3: Phonotactic properties of stimuli). We have also included a detailed report of linguistic features for the target answers in the supplementary file (Table S4: Phonotactic properties of stimuli (cont.)). These features were extracted using the Irvine Phonotactic Online Dictionary (IPhOD) [74]. The auditory naming stimuli were selected from the normative dataset by Hamberger and Seidel [32]. For words not in the list, we created a matched set (see Table S5: Auditory naming stimuli). Participants spontaneously produced the answers without cueing or waiting. The stimuli were randomly interspersed within the block. Visual naming, visual word repetition, and auditory word repetition are repeated twice.

Visual stimuli for visual naming and visual word reading were presented using a laptop (15.6” Retina screen) placed in front of participants at a comfortable distance (0.5m – 1.0m). Auditory stimuli were presented via a speaker placed in front of the presentation laptop. Onsets and offsets of stimuli were detected via analog channels for time-sync purpose: photodiode for visual stimuli and trigger for auditory. Participants responded by speaking into a microphone, with verbal response times extracted from an analog microphone channel.

#### Data Acquisition and Preprocessing

Neural signals were recorded at 2048 Hz with the Nicolet system and were decimated to 512 Hz during export. Electrodes were inspected by epileptologists. Electrodes in the seizure onset zone, with epileptiform activity, or line artifacts were removed from further analysis. The data were then referenced to a common average by averaging the clean signal across all electrodes and subtracting the common average signal from each electrode. Continuous data was divided into epochs based on the onset of stimulus (lock to stimulus onset) or onset of speech (locked to articulation). Articulation onsets were corrected under manual inspection of the audio recordings that are time-synced with the neural recordings. Trials in which participants did not respond or the reaction time was *>*3 standard deviations above the mean reaction time over all trials within each task were removed from analysis as bad trials.

Our analysis of the electrophysiology signals focused on changes in analytic amplitude signal (70–150 Hz). To quantify changes in the high gamma range, data were bandpassed with 8 bands of logarithmically distributed subbands from 70 and 150 Hz and averaged. This multi-band extraction method was used as it avoided the dominance of lower frequency [75, 76]. Analytic amplitude was calculated by taking the absolute value of the Hilbert transform of the filtered signal. The data were then normalized into percent change from baseline with [-250ms, -50]ms prestimulus interval.

### QUANTIFICATION AND STATISTICAL ANALYSIS

#### Task-active Electrode Selection

For each electrode and each task, we computed the analytic amplitude for three cognitive stages: perception (0ms - 500ms locked to stimulus), pre-articulatory (-500ms - 0ms locked to articulation), and articulation (0ms - 500ms locked to articulation). An electrode is considered active if both the maximum value of the mean across trials for any task or period is greater than 50%, and the trial activity is significantly above zero for over 100 ms for either task by computing t-tests across trials against zero (See Supplementary Figure S5 for the number and ratio of task-active electrodes in each region of interest). We took the union of task-active electrodes across tasks and stages for further analysis.

#### Multilinear Factor Model

We fit a multilinear model to test the factors predicting neural activity of individual electrodes. This multilinear encoding model for each electrode predicts neural activity based on the main and interaction effects of the features (fitlm function, Matlab). For each electrode *n*, the activity *y* is modeled as

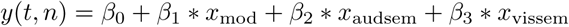

where *y* is a vector with the length of total number of trials denoting the mean high gamma activity over the current time window *t*. *x* is the feature given to the property of the tasks, and *β*s are the weights of each feature being estimated. *x*_mod_ is the binary modality feature that encodes modality, where acoustic is denoted by 1 and visual denoted by 0 (VN, VWR = 0, and AN, AWR = 1). *x*_audsem_ is the binary semantic feature encoding auditory semantics (AN = 1, VN, VWR, AWR = 0, see Fig. 2A). *x*_vissem_ is the binary semantic feature encoding visual semantics (VN = 1, AN, VWR, and AWR = 0). For analysis in Fig. 2B, we chose the time window -750 ms to -250 ms locked to articulation. For analysis in Fig. 2C, we employed a moving window of 250 ms with a hop of 250 ms. The model returns *t* and *p*-values for each *β*. We FDR corrected *p*-value by Benjamini and Hochberg method (mafdr function, Matlab). Electrodes are considered significant using FDR with *q* = 0.001.

#### Non-negative Matrix Factorization

We used non-negative matrix factorization (NMF, nnmf function, Matlab) as a soft clustering technique to cluster the neural data. The data matrix *A* is arranged in the shape of time by electrodes, where the time is composed of concatenated neural activity in perception period (0ms - 500ms), pre-articulatory period (-500ms - 0ms), and articulation (0ms - 500ms) across trials and tasks. Data were downsampled from 512 Hz to 125 Hz (after high gamma extraction) considering computational efficiency before concatenation. Since the analytic amplitude of high gamma computed with multiband extraction is a low-frequency signal, downsampling does not affect the spectra.

### NMF algorithm

Given a nonnegative matrix *A* with shape *u* × *v*, we found nonnegative factor matrices *W* (*u* × *k*) and *H* (*k* × *v*) such that *A* ≈ *WH*. NMF decomposes the data matrix into two matrices, W and H, by minimizing the cost function. We determined the optimal number of clusters *k* by identifying the elbow point (i.e. the point where the rate of variance (*R*^2^) decreases sharply levels off. See Supplementary Fig. S2A, Supplementary Fig. S3A). Variance is defined as *R*^2^ = 1 − *SSE_residual_/SSE_total_*, where Sum of Squared Errors (SSE) is calculated with 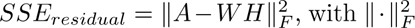 denotes the matrix Frobenius norm that calculates the sum of squared error. Total SSE is defined as 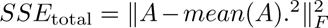. Then, we assigned each electrode to the cluster of its maximum contribution for visualization and future analysis (Supplementary Fig. S2B).

#### Encoding Model and Variance Partitioning

We employed the multivariate temporal response function (mTRF) to relate continuous neural signals and functional stimuli [77, 78]. The continuous neural recordings of the auditory naming task was downsampled (after high gamma extraction) to 125 Hz for computational efficiency. For each electrode, the neural activity is represented as a vector with length *T*, the entire duration from the start of the first trial to the end of the last trial. Three models were constructed to represent different aspects of information Fig. 5A: **Acoustic model:** The acoustic model is constructed by calculating the average across frequency bands of the spectrogram of audio recording for each time point and downsampled to 125 Hz, capturing the acoustic envelope. **Task engagement model:** The task engagement model is a binary model constructed based on the stimulus onset and response offset. For each trial, the model marks one for stimulus onset to response offset, and zero from response offset to next stimulus onset. **Semantic integration model:** We use word surprisal as a measure for semantic load to predict neural processing of higher-level linguistic information in speech [79–82]. Word surprisal is mathematically defined as the negative log probability of a word given its preceding context: *surprisal*(*w_i_*) = − log *P* (*w_i_*|*w*_1_*… w_i−_*_1_). This metric quantifies the relative unexpectedness of a word and reflects the information complexity at each word in a sentence. Here, we constructed the semantic integration model by calculating the increasing surprisal of each word added to the sentence with GPT-2 model [83]. For a sentence with *n* words, the stepwise increment of surprisal is calculated by first getting *n* surprisal values from calculating the first 1, first 1 and 2 words, until 1 to *n* words. Each model is a vector with length *T*, where *T* is the duration of auditory naming task, with a sampling rate of 125 Hz.

The input of three models was normalized to obtain the same magnitude ranging from [0, 1 avoid weighting bias. The mTRF toolbox [77] was employed to estimate *R*^2^ for each model. We used different combinations of the three models to calculate individual contributions.

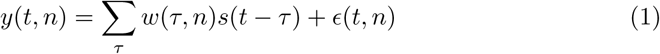

We assumed the output of the system is related to the input via linear convolution. For every electrode, the output is predicted by *n* features, and the instantaneous neural response *y*(*t, n*), sampled at times *t* = 1, · · · *, T*, consists of a convolution of the stimulus property, *s*(*t*), with an unknown electrode-specific temporal response function (TRF), *w*(*τ, n*). *ɛ* is the residual response at each electrode not explained by the model. The range for the time lag was set from 0 ms to 400 ms. To prevent overfitting, we used ridge regression with L2 regularization with 4-fold cross-validation and optimized hyperparameters to maximize the mutual information between actual and predicted responses. This encoding model was fit for all task-active electrodes.

Variance partitioning was performed to separate the unique contribution of each model. To quantify the contribution of different stimulus features to neural activity, we estimated the variance explained (*R*^2^) uniquely by each model and the variance explained by intersections of various combinations of these models (see a more detailed example in Supplementary Fig. S4A and S4B). The *R*^2^ of the sole contribution from one model could be estimated. Set theory was then employed to calculate common (as a set intersection) and unique (as a set difference) variances explained [66, 69].

The statistical significance of the encoding model was assessed using a permutation test. We randomized the continuous neural data by dividing it into 10-second bins, each containing temporal segments with comprehensible acoustic information, and shuffled the order of these bins. This procedure was repeated 10,000 times, and the variance at the 95^th^ percentile was used as the threshold for significance. See Supplementary Table S2 for mean and SEM of the encoding model after variance partitioning in major anatomical regions.

#### Statistical Tests

Unpaired one-way t-test (Fig. 1B) was used to compare whether the neural activity is different from naming tasks and their controls. We first calculated the average neural activity within the pre-articulatory window (-500ms - 0ms) for each trial. The values were compared between naming tasks and their control tasks (i.e. VN vs. VWR; AN vs. AWR) across regions of interest (ttest function, Matlab).

Kruskal-Wallis test (Fig. 3B) was conducted to test whether the distribution of neural activity is different across tasks. We first averaged the neural activity for each electrode across trials within the pre-articulatory window (-500ms - 0ms). For each cluster, Kruskal-Wallis test (kruskalwallis function, Matlab) was used to determine whether data in each task comes from the same distribution. If significant, post hoc analysis was done to determine which pair drives the effect.

Wilcoxon rank sum test (Fig. 5D) was computed to test whether the distribution of unique variance is different between two functional networks (signrank function, Matlab).

#### Bootstrapping and Permutation for Laterality

To evaluate the laterality of word retrieval, we implemented bootstrapping (bootci function, Matlab) to obtain the confidence interval (CI) of the ratios of task-active electrodes in left and right hemispheres. Bootci computes a 95% CI of ratios of task-active electrodes for each hemisphere and cluster, with the number of bootstrap samples set to 10000.

The statistical significance of lateralization was assessed using a permutation test. We hypothesized an equal active ratio in the left and right hemispheres. For each cluster, we first calculated the active ratio of electrodes in both hemispheres by dividing the number of electrodes belonging to the specified cluster by the total number of electrodes in each hemisphere. For permutation, the task-active electrodes in both hemispheres was combined and shuffled to obtain a random distribution of task-active electrodes in total population of both hemispheres. Then, vectors representing left and right hemispheres are split and the new active ratio was calculated. This procedure was repeated 10,000 times. We compared the distribution with the actual active ratio by calculating the *p*-value (i.e. the percentile of true value in the permutation results).

## KEY RESOURCES TABLE

## Acknowledgments

This work was supported by National Institute of Health grants R01NS109367, R01NS115929, and R01DC018805 (A.F.).

## Author contributions

L.Y.: Corresponding author and lead contact; P.D.: Resources; W.D.: Resources; O.D.: Resources; D.F.: Resources; and A.F.: Conceptualization, Funding acquisition, Methodology, Resources, Supervision, and Writing – review & editing.

## Data and Code availability

The data set generated during the current study will be made available from the authors upon request and documentation is provided that the data will be strictly used for research purposes and will comply with the terms of our study IRB. The code is available upon publication at https://github.com/flinkerlab/.

## Declarations

The authors declare that they have no competing interests.

## Supplementary Information

**Fig. S1.**
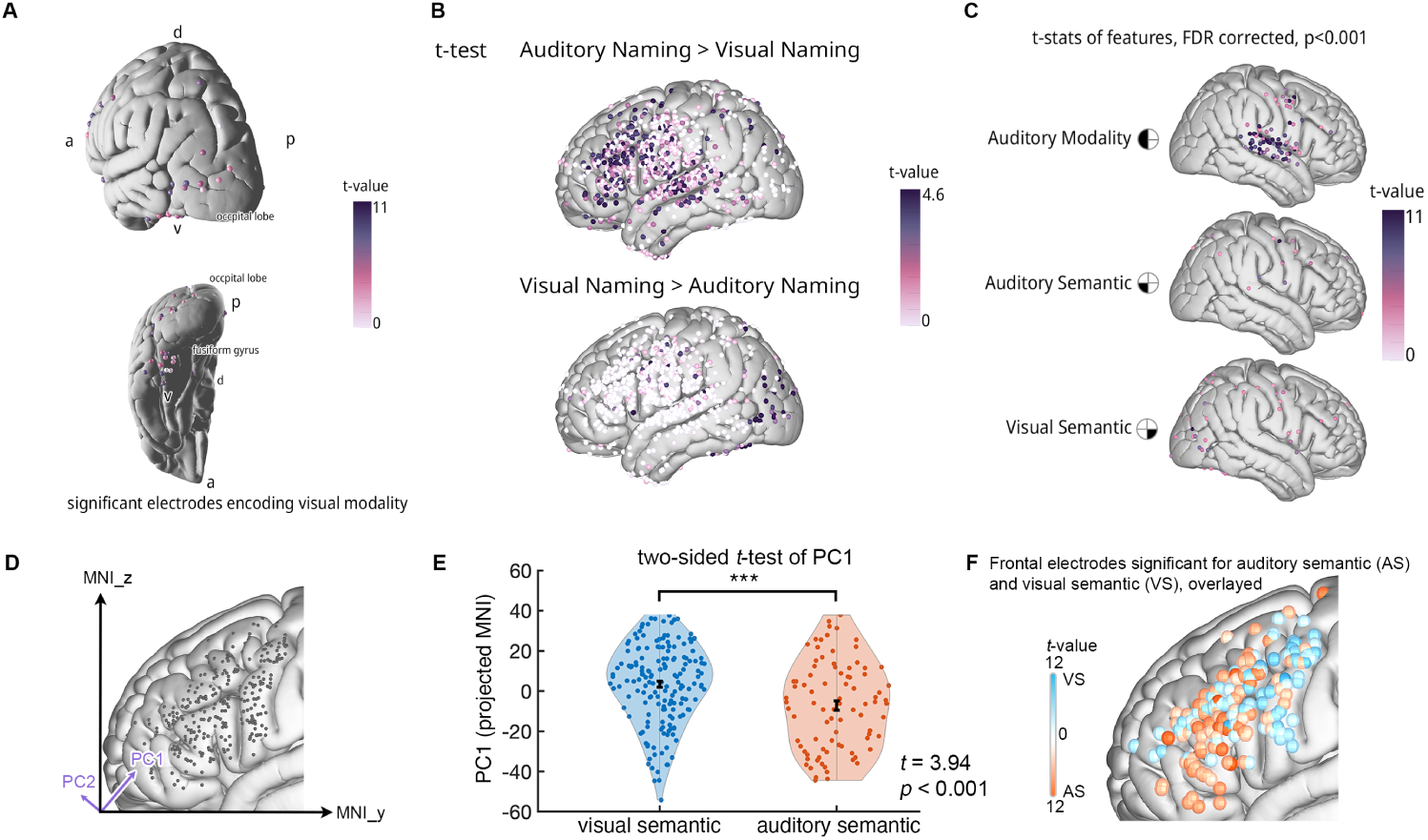
(A) *t*-value map of encoding results -500 ms before articulation. Electrodes that significantly encode the visual modality are projected on the brain (compare with the auditory modality in Fig. 2B, top) (B) *t*-value map of the one-sided *t*-test between auditory naming and visual naming. Significant electrodes are projected on the brain. Unlike the multilinear model in Fig. 2B, a simple *t*-test comparison failed to reveal modality-specific semantic modulation. (C) *t*-value map of encoding results -500 ms before articulation for right hemisphere coverage. Electrodes that significantly encode modality (top), auditory-specific semantics (middle), and visual-specific semantics (bottom) are projected on the brain. (D) We performed a principal component analysis (PCA) on the *MNIy* and *MNIz* coordinates of all task-active frontal electrodes (N=414) to find the principal component that captures the largest explained variance (75.5%). This PC (PC1, in purple) features an axis positioned ventral anterior to dorsal posterior across frontal cortex. (E)) We compared the projected spatial coordinates (PC values) with a two-sample *t*-test between the visual semantic (N=170) and auditory semantic (N=87) groups. The values are visualized in violin plots, and error bars represent the group mean and SEM. Visual semantic electrodes were significantly larger, indicating the group was located more dorsal posteriorly. (F) Visualization of electrodes significantly encoding either auditory or visual semantics, overlayed on the same cortical surface.

**Fig. S2.**
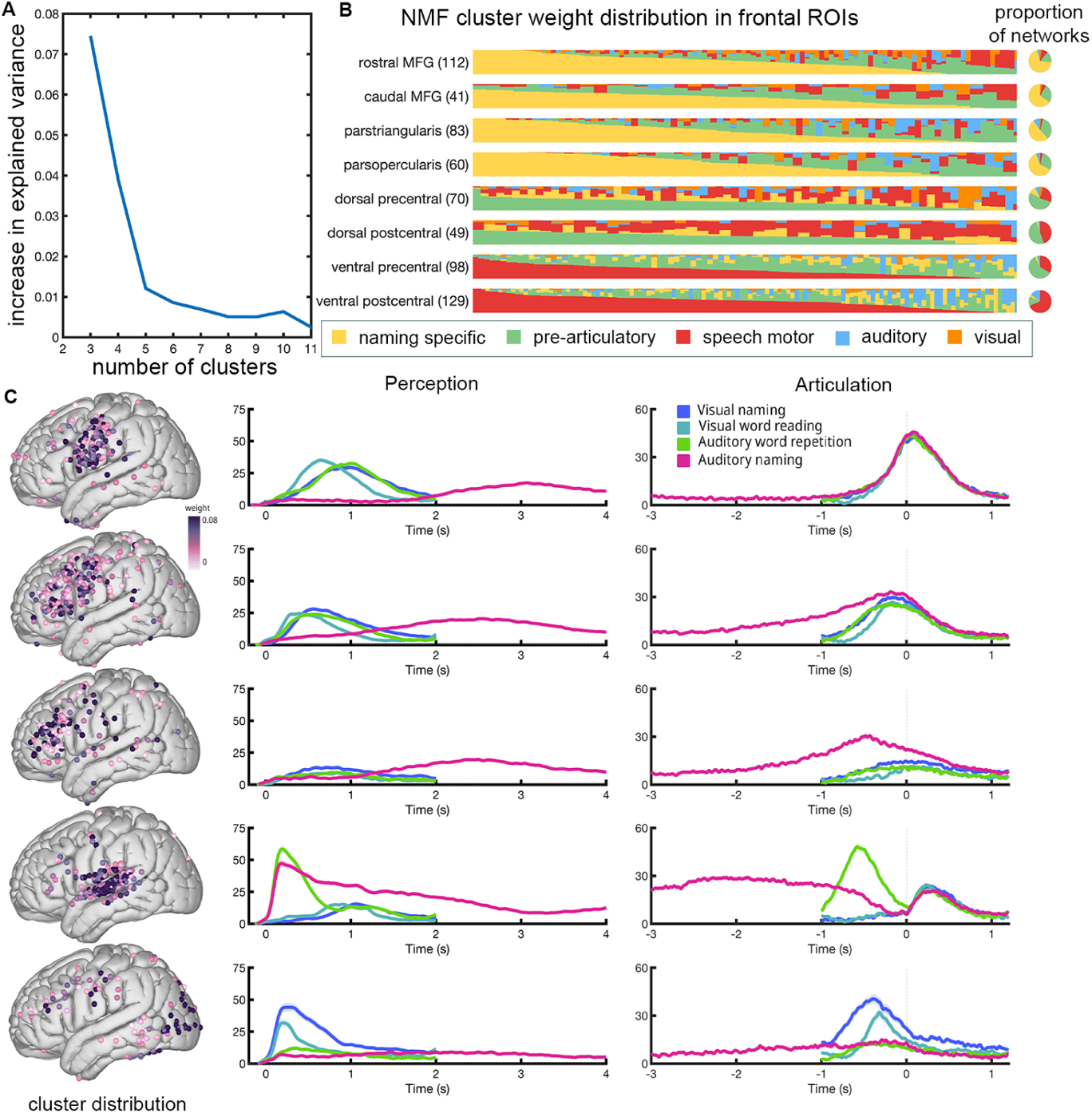
Clustering results in a cohort with left hemisphere coverage. (A) The number of optimal clusters was determined based on the elbow point. (B) Left: response pattern of individual electrodes across each ROI. The numbers indicate the total electrodes within each ROI. Each column represents one electrode with the distribution of weights assigned per cluster, normalized so that the weights sum to 1. Each electrode is eventually assigned a “winning” cluster representing the maximal contribution (depicted as colored area in the column). Electrodes within each ROI are sorted based on dominant clusters for that ROI. Right: pie chart showing the proportion of final “winning” clusters for each ROI. Dorsal precentral gyrus contains five networks, and has the highest ratio (22%) of combined sensory and naming-specific clusters, compared to the other subdivisions in speech motor cortex. (C) Left: the NMF weight distribution of electrodes in all clusters. Right: the weighted average of neural high gamma activity across electrodes in the cluster, locked to perception and articulation.

**Fig. S3.**
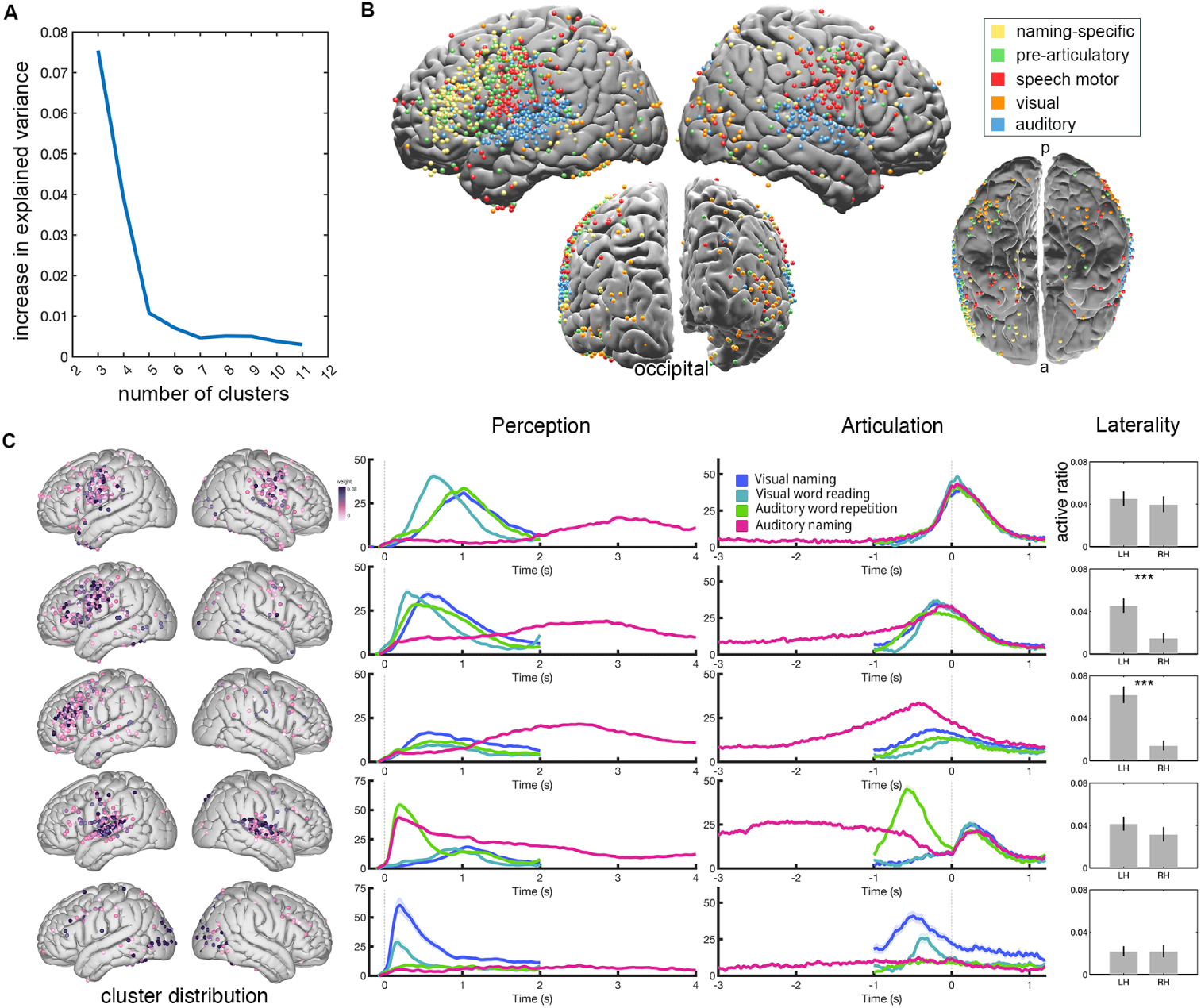
Clustering results in a cohort with bilateral hemisphere coverage. (A) The number of optimal clusters was determined based on the elbow point. (B) Spatial positions of electrodes color-coded by NMF cluster membership. (C) Left: the NMF weight distribution of electrodes in all clusters. Middle: the weighted average of neural high gamma activity across electrodes in the cluster, locked to perception and articulation. Right: active ratios (task-active electrodes vs. all) for the left and right hemispheres are shown with bootstrapped confidence intervals depicted in the error bar. Significance between left and right hemisphere active ratio was determined by a permutation test.

**Fig. S4.**
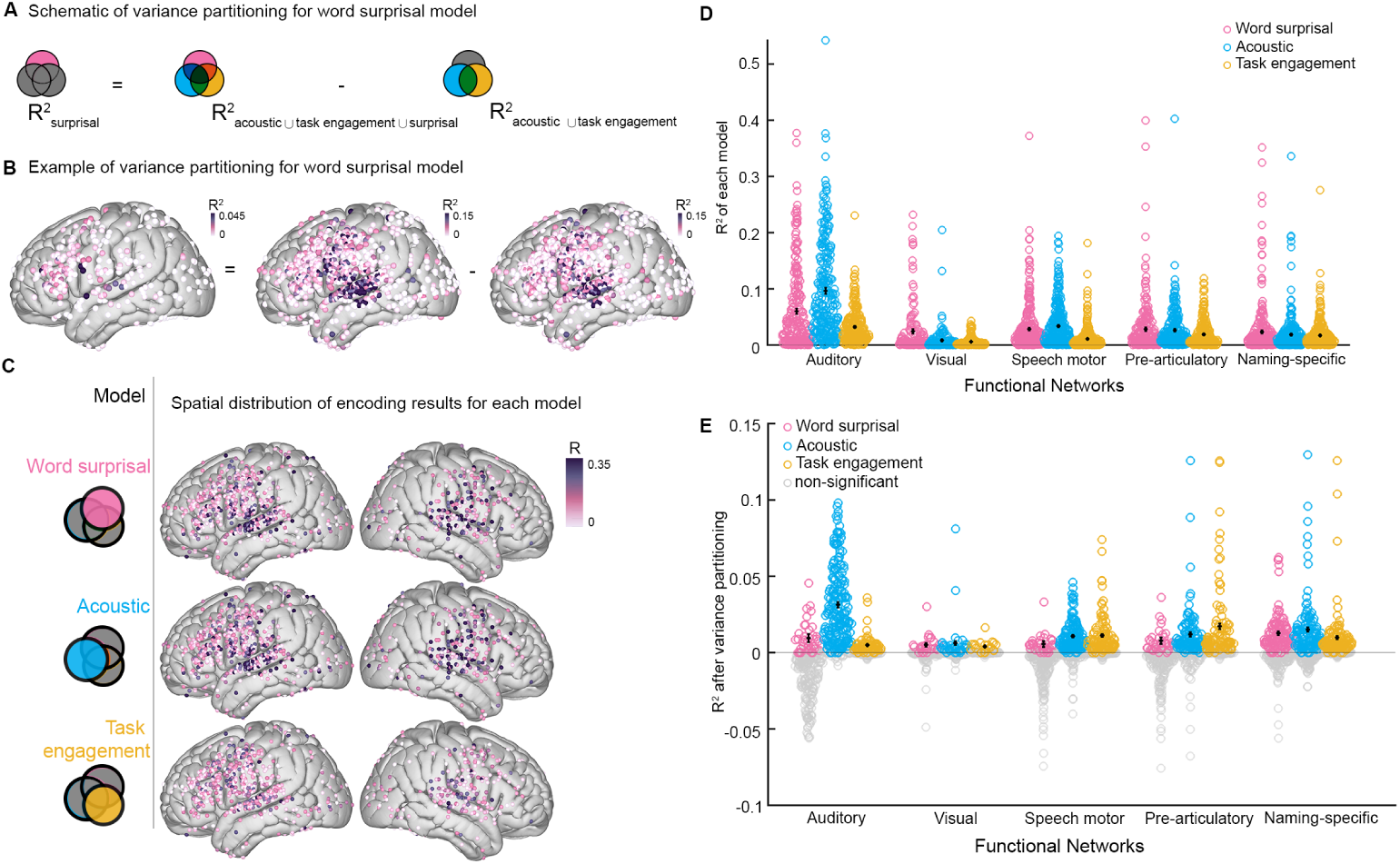
(A) Exemplar schematic of variance partitioning for word surprisal model. (B) Visualization of the calculation of explained R^2^ across electrodes. (C) R^2^ of individual mTRF model across electrodes. (D) Distribution of R^2^ by each mTRF model within each network prior to variance partitioning. (E) Distribution of unique variance explained by each mTRF model within each network identified by NMF. Significant models are shown in color (permutation test).

**Fig. S5.**
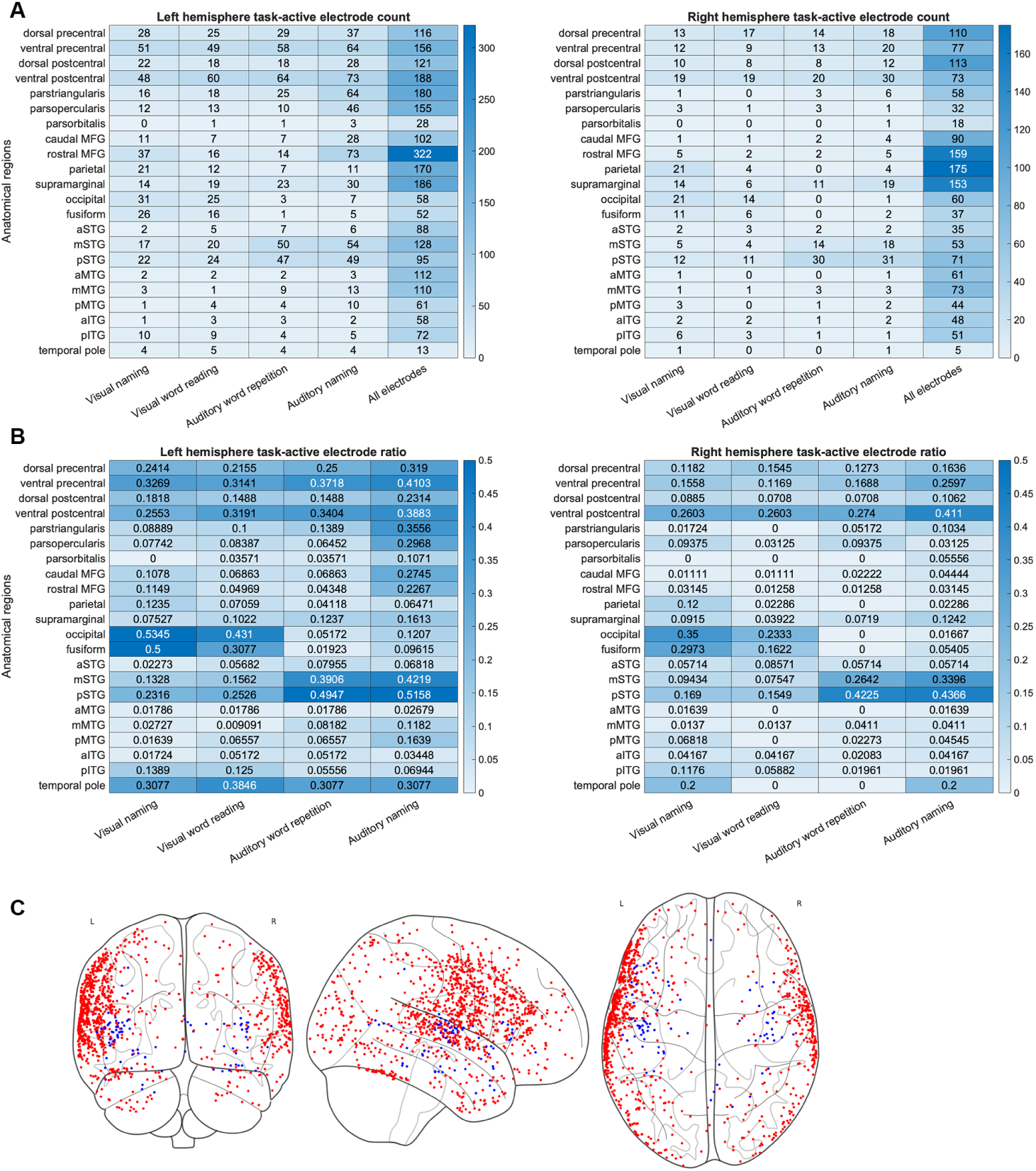
(A) Task-active electrode and total electrode count in major anatomical regions. (B) Active ratio for in major anatomical regions. (C) Electrode coverage distribution. Surface electrodes are marked in red, and depth electrodes are marked in blue.

**Table S1.** Participant Demographics.

| ID | Coverage | Gender | Handedness | Age at Surgery | VCI | Wada Language |
| --- | --- | --- | --- | --- | --- | --- |
| 1 | Right | Male | Right | 28 | 134 | - |
| 2 | Right | Male | Right | 22 | 114 | Left |
| 3 | Right | Male | Left | 26 | - | - |
| 4 | Bilateral | Male | Right | 19 | 105 | Left |
| 5 | Left | Female | Right | 55 | 102 | Left |
| 6 | Left | Female | Right | 41 | 126 | - |
| 7 | Right | Male | Left | 30 | 116 | - |
| 8 | Left | Female | Right | 49 | 107 | Left |
| 9 | Right | Male | Left | 47 | 138 | - |
| 10 | Left | Male | Right | 18 | 95 | - |
| 11 | Bilateral | Male | Right | 26 | - | Left |
| 12 | Left | Female | Right | 20 | 118 | Left |
| 13 | Right | Male | Right | 16 | 108 | - |
| 14 | Right | Female | Right | 21 | 95 | Left |
| 15 | Right | Male | - | 10 | 123 | - |
| 16 | Left | Female | Right | 34 | - | Left |
| 17 | Left | Female | Right | 30 | 100 | Left |
| 18 | Right | Male | Left | 36 | 116 | Left |
| 19 | Right | Female | Right | 26 | 70 | - |
| 20 | Left | Female | - | 56 | 127 | - |
| 21 | Left | Male | Right | 22 | 96 | Left |
| 22 | Bilateral | Male | Right | 57 | 91 | Left |
| 23 | Right | Male | Right | 37 | 110 | - |
| 24 | Left | Female | Left | 23 | 91 | Left |
| 25 | Left | Male | Right | 28 | - | - |
| 26 | Left | Male | Right | 43 | 87 | Left |
| 27 | Left | Male | Right | 41 | 103 | - |
| 28 | Left | Male | Right | 47 | 130 | Left |
| 29 | Left | Male | Right | 18 | 98 | Left |
| 30 | Right | Female | Left | 27 | 96 | Left |
| 31 | Left | Female | Right | 33 | 70 | - |
| 32 | Left | Female | Right | 30 | 98 | Left |
| 33 | Left | Female | Right | 28 | 102 | Bilateral |
| 34 | Left | Male | Right | 24 | 145 | Left |
| 35 | Left | Female | Right | 23 | 79 | Left |
| 36 | Right | Male | Right | 63 | 96 | - |
| 37 | Left | Female | Left | 14 | 81 | - |
| 38 | Right | Female | Right | 49 | 91 | Left |
| 39 | Left | Male | Right | 31 | 116 | Left |
| 40 | Right | Male | Right | 17 | - | - |
| 41 | Stereo | Male | Right | 14 | 92 | - |
| 42 | Right | Female | Right | 31 | 104 | - |
| 43 | Left | Male | Right | 20 | 95 | Left |
| 44 | Right | Female | Left | 30 | 107 | Left |
| 45 | Bilateral | Male | Right | 20 | 85 | - |
| 46 | Right | Female | Right | 38 | 100 | Left |
| 47 | Left | Male | Right | 25 | 76 | Left |
| 48 | Left | Male | Right | 37 | 102 | Left |

**Table S2.**
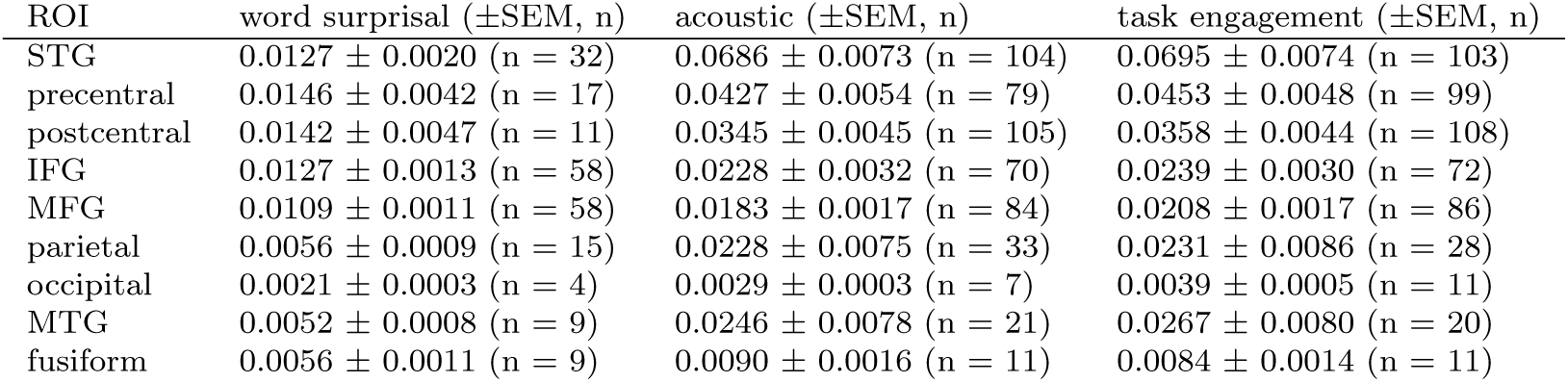
Mean and SEM of the encoding model after variance partitioning per ROI.

| ROI | word surprisal (±SEM, n) | acoustic (±SEM, n) | task engagement (±SEM, n) |
| --- | --- | --- | --- |
| STG | 0.0127 ± 0.0020 (n = 32) | 0.0686 ± 0.0073 (n = 104) | 0.0695 ± 0.0074 (n = 103) |
| precentral | 0.0146 ± 0.0042 (n = 17) | 0.0427 ± 0.0054 (n = 79) | 0.0453 ± 0.0048 (n = 99) |
| postcentral | 0.0142 ± 0.0047 (n = 11) | 0.0345 ± 0.0045 (n = 105) | 0.0358 ± 0.0044 (n = 108) |
| IFG | 0.0127 ± 0.0013 (n = 58) | 0.0228 ± 0.0032 (n = 70) | 0.0239 ± 0.0030 (n = 72) |
| MFG | 0.0109 ± 0.0011 (n = 58) | 0.0183 ± 0.0017 (n = 84) | 0.0208 ± 0.0017 (n = 86) |
| parietal | 0.0056 ± 0.0009 (n = 15) | 0.0228 ± 0.0075 (n = 33) | 0.0231 ± 0.0086 (n = 28) |
| occipital | 0.0021 ± 0.0003 (n = 4) | 0.0029 ± 0.0003 (n = 7) | 0.0039 ± 0.0005 (n = 11) |
| MTG | 0.0052 ± 0.0008 (n = 9) | 0.0246 ± 0.0078 (n = 21) | 0.0267 ± 0.0080 (n = 20) |
| fusiform | 0.0056 ± 0.0011 (n = 9) | 0.0090 ± 0.0016 (n = 11) | 0.0084 ± 0.0014 (n = 11) |

**Table S3.**
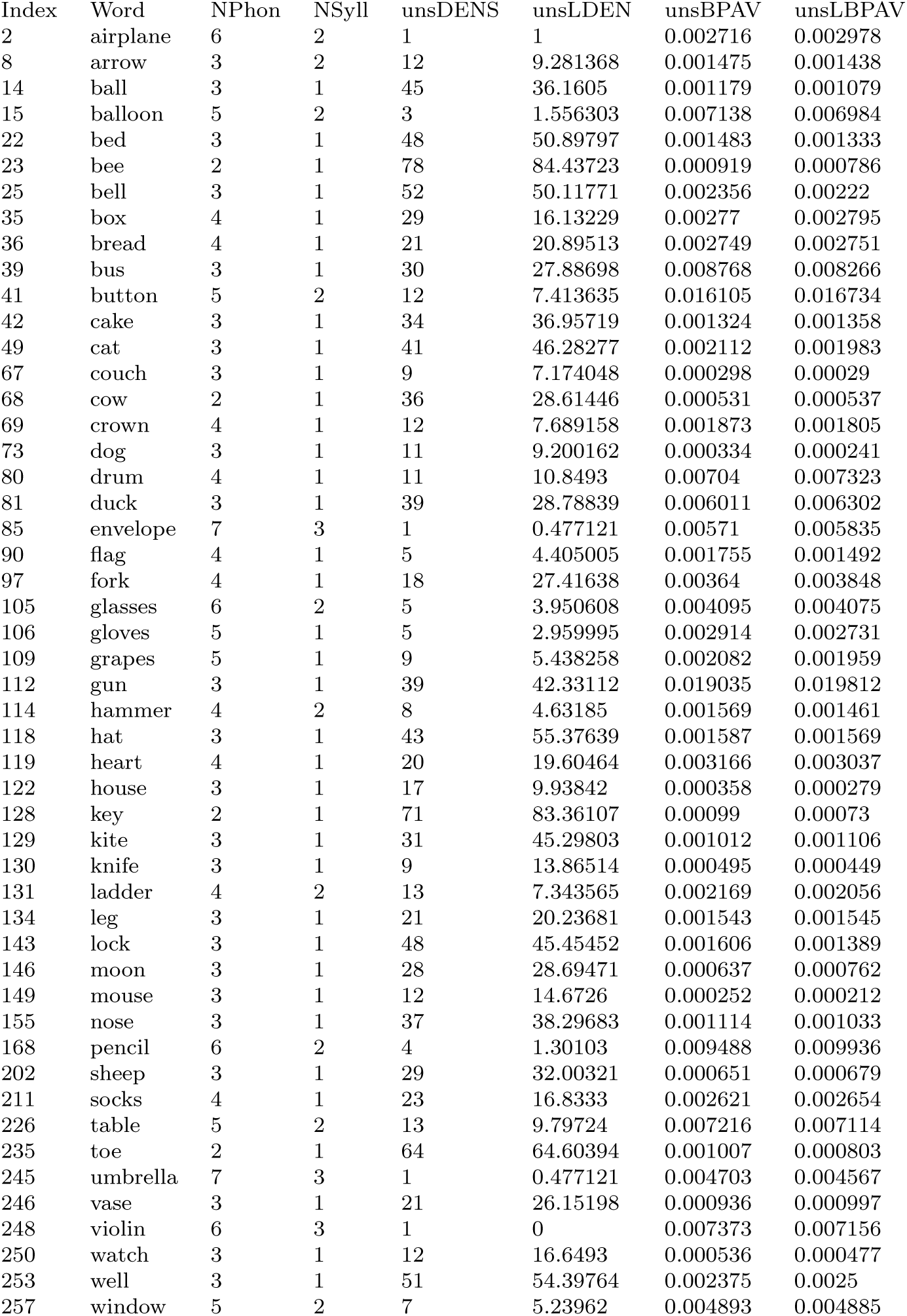
Phonotactic properties of stimuli. Index is matched with visual naming dataset index. NPhon: number of phonemes. NSyll: number of syllables. unsDENS: unstressed phonological neighborhood density; vowel-stress ignored. unsLDEN: unsDENS weighted with Kucera-Francis log frequency of neighbors. unsBPAV: stressed biphoneme probability average; distinct stressed-vowels. unsLBPAV: unsBPAV weighted with log Kucera-Francis word frequency.

| Index | Word | NPhon | NSyll | unsDENS | unsLDEN | unsBPAV | unsLBPAV |
| --- | --- | --- | --- | --- | --- | --- | --- |
| 2 | airplane | 6 | 2 | 1 | 1 | 0.002716 | 0.002978 |
| 8 | arrow | 3 | 2 | 12 | 9.281368 | 0.001475 | 0.001438 |
| 14 | ball | 3 | 1 | 45 | 36.1605 | 0.001179 | 0.001079 |
| 15 | balloon | 5 | 2 | 3 | 1.556303 | 0.007138 | 0.006984 |
| 22 | bed | 3 | 1 | 48 | 50.89797 | 0.001483 | 0.001333 |
| 23 | bee | 2 | 1 | 78 | 84.43723 | 0.000919 | 0.000786 |
| 25 | bell | 3 | 1 | 52 | 50.11771 | 0.002356 | 0.00222 |
| 35 | box | 4 | 1 | 29 | 16.13229 | 0.00277 | 0.002795 |
| 36 | bread | 4 | 1 | 21 | 20.89513 | 0.002749 | 0.002751 |
| 39 | bus | 3 | 1 | 30 | 27.88698 | 0.008768 | 0.008266 |
| 41 | button | 5 | 2 | 12 | 7.413635 | 0.016105 | 0.016734 |
| 42 | cake | 3 | 1 | 34 | 36.95719 | 0.001324 | 0.001358 |
| 49 | cat | 3 | 1 | 41 | 46.28277 | 0.002112 | 0.001983 |
| 67 | couch | 3 | 1 | 9 | 7.174048 | 0.000298 | 0.00029 |
| 68 | cow | 2 | 1 | 36 | 28.61446 | 0.000531 | 0.000537 |
| 69 | crown | 4 | 1 | 12 | 7.689158 | 0.001873 | 0.001805 |
| 73 | dog | 3 | 1 | 11 | 9.200162 | 0.000334 | 0.000241 |
| 80 | drum | 4 | 1 | 11 | 10.8493 | 0.00704 | 0.007323 |
| 81 | duck | 3 | 1 | 39 | 28.78839 | 0.006011 | 0.006302 |
| 85 | envelope | 7 | 3 | 1 | 0.477121 | 0.00571 | 0.005835 |
| 90 | flag | 4 | 1 | 5 | 4.405005 | 0.001755 | 0.001492 |
| 97 | fork | 4 | 1 | 18 | 27.41638 | 0.00364 | 0.003848 |
| 105 | glasses | 6 | 2 | 5 | 3.950608 | 0.004095 | 0.004075 |
| 106 | gloves | 5 | 1 | 5 | 2.959995 | 0.002914 | 0.002731 |
| 109 | grapes | 5 | 1 | 9 | 5.438258 | 0.002082 | 0.001959 |
| 112 | gun | 3 | 1 | 39 | 42.33112 | 0.019035 | 0.019812 |
| 114 | hammer | 4 | 2 | 8 | 4.63185 | 0.001569 | 0.001461 |
| 118 | hat | 3 | 1 | 43 | 55.37639 | 0.001587 | 0.001569 |
| 119 | heart | 4 | 1 | 20 | 19.60464 | 0.003166 | 0.003037 |
| 122 | house | 3 | 1 | 17 | 9.93842 | 0.000358 | 0.000279 |
| 128 | key | 2 | 1 | 71 | 83.36107 | 0.00099 | 0.00073 |
| 129 | kite | 3 | 1 | 31 | 45.29803 | 0.001012 | 0.001106 |
| 130 | knife | 3 | 1 | 9 | 13.86514 | 0.000495 | 0.000449 |
| 131 | ladder | 4 | 2 | 13 | 7.343565 | 0.002169 | 0.002056 |
| 134 | leg | 3 | 1 | 21 | 20.23681 | 0.001543 | 0.001545 |
| 143 | lock | 3 | 1 | 48 | 45.45452 | 0.001606 | 0.001389 |
| 146 | moon | 3 | 1 | 28 | 28.69471 | 0.000637 | 0.000762 |
| 149 | mouse | 3 | 1 | 12 | 14.6726 | 0.000252 | 0.000212 |
| 155 | nose | 3 | 1 | 37 | 38.29683 | 0.001114 | 0.001033 |
| 168 | pencil | 6 | 2 | 4 | 1.30103 | 0.009488 | 0.009936 |
| 202 | sheep | 3 | 1 | 29 | 32.00321 | 0.000651 | 0.000679 |
| 211 | socks | 4 | 1 | 23 | 16.8333 | 0.002621 | 0.002654 |
| 226 | table | 5 | 2 | 13 | 9.79724 | 0.007216 | 0.007114 |
| 235 | toe | 2 | 1 | 64 | 64.60394 | 0.001007 | 0.000803 |
| 245 | umbrella | 7 | 3 | 1 | 0.477121 | 0.004703 | 0.004567 |
| 246 | vase | 3 | 1 | 21 | 26.15198 | 0.000936 | 0.000997 |
| 248 | violin | 6 | 3 | 1 | 0 | 0.007373 | 0.007156 |
| 250 | watch | 3 | 1 | 12 | 16.6493 | 0.000536 | 0.000477 |
| 253 | well | 3 | 1 | 51 | 54.39764 | 0.002375 | 0.0025 |
| 257 | window | 5 | 2 | 7 | 5.23962 | 0.004893 | 0.004885 |

**Table S4.** Phonotactic properties of stimuli (cont.) unsPOSPAV: unstressed positional probability; vowel-stress ignored. unsLPOSPAV: unsPOSPAV weighted with log Kucera-Francis frequency. KFFREQ: Kucera-Francis Written Word Frequency for real words. LOGFRQ: log Kucera-Francis Written Word Frequency for real words.

| Word | unsPOSPAV | unsLPOSPAV | KFFREQ | LOGFRQ |
| --- | --- | --- | --- | --- |
| airplane | 0.0490955 | 0.0487499 | 11 | 1.04139 |
| arrow | 0.041846 | 0.0415525 | 14 | 1.14613 |
| ball | 0.0575015 | 0.0533727 | 117 | 2.06819 |
| balloon | 0.0657108 | 0.0647619 | 10 | 1 |
| bed | 0.0581851 | 0.0563127 | 135 | 2.13033 |
| bee | 0.050951 | 0.0470635 | 13 | 1.11394 |
| bell | 0.0705603 | 0.0677808 | 18 | 1.25527 |
| box | 0.0599479 | 0.0572366 | 74 | 1.86923 |
| bread | 0.0555971 | 0.0555032 | 41 | 1.61278 |
| bus | 0.0801274 | 0.0798818 | 34 | 1.53148 |
| button | 0.0899062 | 0.0902106 | 12 | 1.07918 |
| cake | 0.0604746 | 0.061174 | 13 | 1.11394 |
| cat | 0.073916 | 0.0727082 | 24 | 1.38021 |
| couch | 0.0375644 | 0.0379081 | 13 | 1.11394 |
| cow | 0.0522283 | 0.0520486 | 31 | 1.49136 |
| crown | 0.0588186 | 0.0586101 | 20 | 1.30103 |
| dog | 0.0379572 | 0.0366465 | 77 | 1.88649 |
| drum | 0.056979 | 0.0560157 | 11 | 1.04139 |
| duck | 0.0731905 | 0.073737 | 9 | 0.954243 |
| envelope | 0.0486021 | 0.0473771 | 21 | 1.32222 |
| flag | 0.0338066 | 0.0325638 | 18 | 1.25527 |
| fork | 0.0509711 | 0.0509317 | 15 | 1.17609 |
| glasses | 0.0612227 | 0.0606609 | 29 | 1.4624 |
| gloves | 0.0364713 | 0.0347361 | 7 | 0.845098 |
| grapes | 0.04589 | 0.0459107 | 7 | 0.845098 |
| gun | 0.0757774 | 0.0773877 | 122 | 2.08636 |
| hammer | 0.053852 | 0.0516623 | 10 | 1 |
| hat | 0.0551715 | 0.0535211 | 57 | 1.75587 |
| heart | 0.0663672 | 0.0671005 | 175 | 2.24304 |
| house | 0.0417534 | 0.0421095 | 601 | 2.77887 |
| key | 0.0677762 | 0.0671264 | 92 | 1.96379 |
| kite | 0.063032 | 0.0643507 | 1 | 0 |
| knife | 0.024967 | 0.0247333 | 80 | 1.90309 |
| ladder | 0.0508569 | 0.0489293 | 20 | 1.30103 |
| leg | 0.0421822 | 0.0422153 | 62 | 1.79239 |
| lock | 0.0518166 | 0.0496278 | 25 | 1.39794 |
| moon | 0.0579849 | 0.0583421 | 66 | 1.81954 |
| mouse | 0.0474466 | 0.0476799 | 10 | 1 |
| nose | 0.0253865 | 0.0268207 | 60 | 1.77815 |
| pencil | 0.0841065 | 0.086527 | 36 | 1.5563 |
| sheep | 0.0297419 | 0.0292073 | 24 | 1.38021 |
| socks | 0.0713367 | 0.0716347 | 7 | 0.845098 |
| table | 0.0610925 | 0.0605179 | 204 | 2.30963 |
| toe | 0.0358729 | 0.0362062 | 12 | 1.07918 |
| umbrella | 0.0519389 | 0.0518569 | 8 | 0.90309 |
| vase | 0.0412988 | 0.0438164 | 4 | 0.60206 |
| violin | 0.0461178 | 0.0462018 | 11 | 1.04139 |
| watch | 0.0333075 | 0.0334552 | 85 | 1.92942 |
| well | 0.0592139 | 0.0600127 | 1004 | 3.00173 |
| window | 0.0536467 | 0.0552205 | 120 | 2.07918 |

**Table S5.** Auditory naming stimuli.

| Number | Description | Answer |
| --- | --- | --- |
| 1 | a vehicle that flies through the air | airplane |
| 2 | an object shot at a target with a bow | arrow |
| 3 | a round object that you kick | ball |
| 4 | a rubber object that you inflate with helium | balloon |
| 5 | a furniture that you sleep on at night | bed |
| 6 | an insect that produces honey | bee |
| 7 | a musical instrument that makes a ringing sound | bell |
| 8 | a cardboard container | box |
| 9 | a popular food made of flour used in sandwiches | bread |
| 10 | what children ride to school in | bus |
| 11 | a coin shaped object that fastens your dress shirt closed | button |
| 12 | a frosted dessert with a birthday candle | cake |
| 13 | a household pet that purrs | cat |
| 14 | a piece of furniture that multiple people sit on | couch |
| 15 | an animal that produces milk | cow |
| 16 | what a king wears on his head | crown |
| 17 | an animal that barks | dog |
| 18 | an instrument you beat with sticks | drum |
| 19 | an animal that quacks | duck |
| 20 | what you put a letter in and a stamp on | envelope |
| 21 | a rectangular cloth that stands for a nation | flag |
| 22 | a dining utensil used to eat salad | fork |
| 23 | a pair of lenses you wear on your face to help you see | glasses |
| 24 | a piece of clothing that you wear on your hands | gloves |
| 25 | a purple cluster of fruits that grow from a vine | grapes |
| 26 | a weapon that shoots bullets | gun |
| 27 | a tool for hitting nails | hammer |
| 28 | a piece of clothing that you wear on your head | hat |
| 29 | the organ that pumps blood | heart |
| 30 | a building that a family lives in | house |
| 31 | the thing that unlocks a door | key |
| 32 | a diamond shaped toy flown in the wind attached to a string | kite |
| 33 | an eating utensil used for slicing | knife |
| 34 | an equipment used for climbing up and down something | ladder |
| 35 | the limbs you used to walk with | leg |
| 36 | a device to secure a door | lock |
| 37 | an object in outer space that revolves around the earth | moon |
| 38 | a tiny rodent that eats cheese | mouse |
| 39 | a part of your face you use to smell | nose |
| 40 | an instrument used for drawing that is erasable | pencil |
| 41 | an adult farm animal with a wool coat | sheep |
| 42 | a garment you put on your feet before your shoes | socks |
| 43 | a piece of furniture that you eat your dinner on | table |
| 44 | one of the five digits on your foot | toe |
| 45 | the hand-held object that keeps you dry in the rain | umbrella |
| 46 | a decorative container to display flowers | vase |
| 47 | a stringed instrument that rests under your chin | violin |
| 48 | a device you wear on your wrist that tells time | watch |
| 49 | a deep hole in the ground where people obtain water | well |
| 50 | openings in the room fitted with glass | window |

